# Formation and role of the portal of *Staphylococcus aureus* bacteriophage 80α

**DOI:** 10.64898/2026.02.04.703111

**Authors:** Amarshi Mukherjee, James L. Kizziah, Laura K. Parker, Patrick M. Lindstrom, Nehaal Akavaram, Terje Dokland

**Author notes:** Communicating author: Terje Dokland, Department of Microbiology, The University of Alabama at Birmingham, Birmingham, AL 35294, United States of America.

## Abstract

Bacteriophages play an important role in the pathogenicity of *Staphylococcus aureus*, an important human pathogen. Phages are involved in generalized and specialized transduction as well as a more specific process by which they mobilize elements known as phage-inducible chromosomal islands, of which *S. aureus* pathogenicity islands (SaPIs) are an important group. SaPIs are mobilized at high frequency through interactions with specific “helper” bacteriophages, such as 80α, leading to packaging of the SaPI genomes into virions made from structural proteins supplied by the helper. Among these structural proteins is the portal protein, which forms a ring-like portal at a fivefold vertex of the capsid, through which the DNA is packaged during virion assembly and ejected upon infection of the host. We previously showed that portal protein expressed in *E. coli* forms tridecameric rings, while portals found in virions are always dodecamers. To understand the role of the portal in capsid assembly, DNA packaging and ejection, we have here examined this phenomenon further. We show that portals assembled at lower temperature form unclosed rings that may represent portal assembly intermediates. By analyzing portal protein deletion mutants, we demonstrate the involvement of the different functional domains in phage assembly and protein incorporation.

## INTRODUCTION

*Staphylococcus aureus* is an opportunistic human pathogen and one of the leading causes of deaths associated with antibiotic resistance [1]. *S. aureus* encodes an arsenal of virulence factors, many of which are encoded on mobile genetic elements (MGEs), such as plasmids, bacteriophages (phages) and chromosomal islands [2–4]. Tailed bacteriophages with double-stranded (ds) DNA genomes belonging to class *Caudoviricetes* play a key role in the evolution of bacterial pathogenicity, especially in species such as *Staphylococcus aureus*, which are not generally transformable and rarely undergo conjugation [5].

Phage 80α is a typical example of a temperate staphylococcal phage, consisting of a 43,864 base pair double-stranded DNA genome [6](NCBI RefSeq NC_009526.1) packaged inside a 63 nm icosahedral head attached to a 190 nm long tail [7–10]. Like other members of the *Caudoviricetes* phages, 80α assembles a precursor capsid (procapsid) from capsid protein (CP; gp47), a scaffolding protein (SP; gp46), and a portal protein (PP; gp42) [11, 12] (Fig. 1A). PP forms a ring-like portal at one vertex of the capsid, which is assumed to serve as the nucleus for capsid assembly, the entry and exit point for the DNA and as attachment point for the tails [13–15]. Protein gp44 is a minor capsid protein (ejection protein, EP) that is ejected together with the DNA during infection and is important for the survival of the phage DNA post injection [16] (Fig. 1B). The phage genome is packaged through the portal by the terminase, consisting of large (TerL) and small (TerS) subunits [17], followed by capsid maturation, which involves shell expansion, removal of the SP, and attachment of the tail (Fig. 1B).

**Figure 1.**
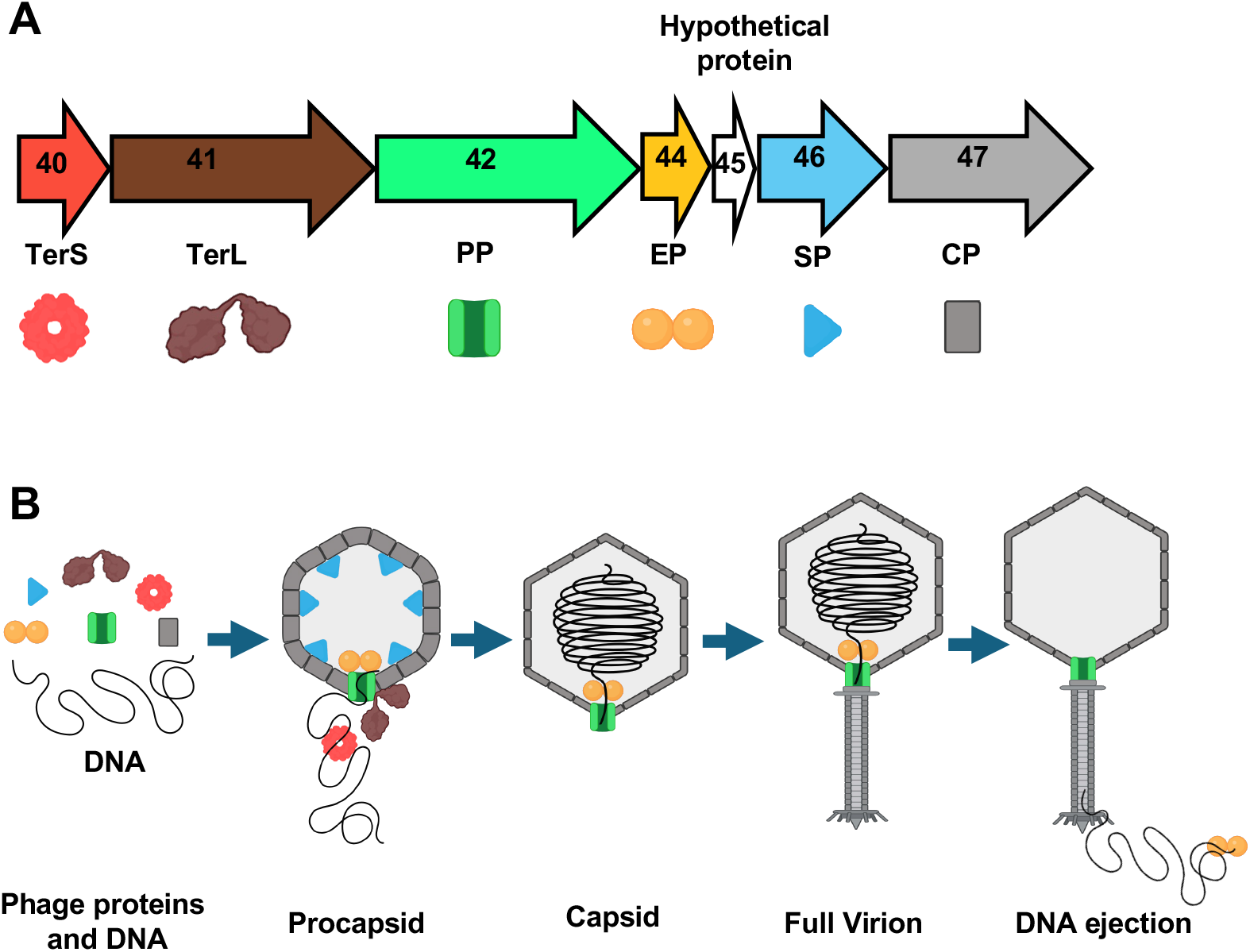
The 80α capsid gene cluster. **(A)** Schematic diagram of ORFs 40–47, including genes encoding small terminase (TerS, gp40), large terminase (TerL, gp41), portal protein (PP, gp42), ejection protein (EP, gp44), scaffolding protein (SP, gp46), and capsid protein (CP, gp47). **(B)** Schematic diagram showing proteins involved in capsid assembly, DNA packaging and DNA ejection.

Phages closely related to 80α are found in a wide variety of pathogenic staphylococcal strains [18]. 80α and its relatives are known as “helper” phages that involved in the specifical mobilization of elements known as phage-inducible chromosomal islands (PICIs) [19], of which the *S. aureus* pathogenicity islands (SaPIs) form an important group. SaPIs encode a range of virulence factors, including superantigen toxins, adhesins and anti-coagulation factors [20, 21]. When SaPIs are induced via derepression by a helper phage [22], the element excises from the host genome, replicates and becomes packaged into transducing particles constructed from phage-encoded structural proteins. In many cases, SaPIs redirect the assembly pathway of its helper phage to form capsids of a smaller size than that normally made by the phage [22, 23], consistent with the smaller size of the SaPI genome [24, 25].

We previously showed that expression of the 80α PP in *Escherichia coli* led to the formation of tridecameric rings, whereas portals found in SaPI1 (and presumably 80α) virions were dodecameric [26], raising the question of how dodecameric portals are formed and incorporated into capsids. What are the regulatory mechanisms that control the oligomeric state of the portal? Here, we have addressed these questions by examining assemblies made by PP expressed under different conditions and with mutations in key domains predicted to interact with other phage components. Our results suggest that portals are initially formed as unstable, unclosed rings, which can be converted into closed dodecamers through interaction with other phage proteins, whereas in solution, the tridecamer is the thermodynamically more stable conformation. The PP clip domain is especially important in this process, implicating the terminase and/or neck proteins in the assembly process. We also showed that the PP crown domain is involved in incorporation of the gp44 ejection protein (EP). The N-terminal wing domain is required for the stability of the protein as well as capsid assembly. Our results underscore the central role of the portal in the assembly and DNA packaging in 80α and other phages, and highlights the complex dynamics that underpin this process.

## RESULTS

### 1. Expression of portal protein in *E. coli* and *S. aureus*

We previously showed that expression of 80α PP (gp42) in *E. coli* led to the formation of tridecameric rings [26]. Since portals found in capsids are invariably dodecamers, the formation of tridecamers appeared to be an artifact of the expression conditions. To address whether the formation of dodecamers was dependent on *S. aureus*-specific host factors, we expressed C-terminally His-tagged PP [26] in *S. aureus* [27] followed by purification by Ni-NTA affinity and size exclusion chromatography (SEC). Both the *E. coli* and *S. aureus*-expressed portals purified in a single peak consistent with portal oligomers (Fig. 2A). The purified portals were imaged by negative stain EM (Fig. 2B), followed by 2D classification in cryoSPARC [28], which showed that the predominant form of both the *E. coli*- and *S. aureus*-produced portals was a tridecamer (Fig. 2C). No dodecameric portals were observed, indicating that the formation of dodecamers was not dependent on *S. aureus*-specific factors alone.

**Figure 2.**
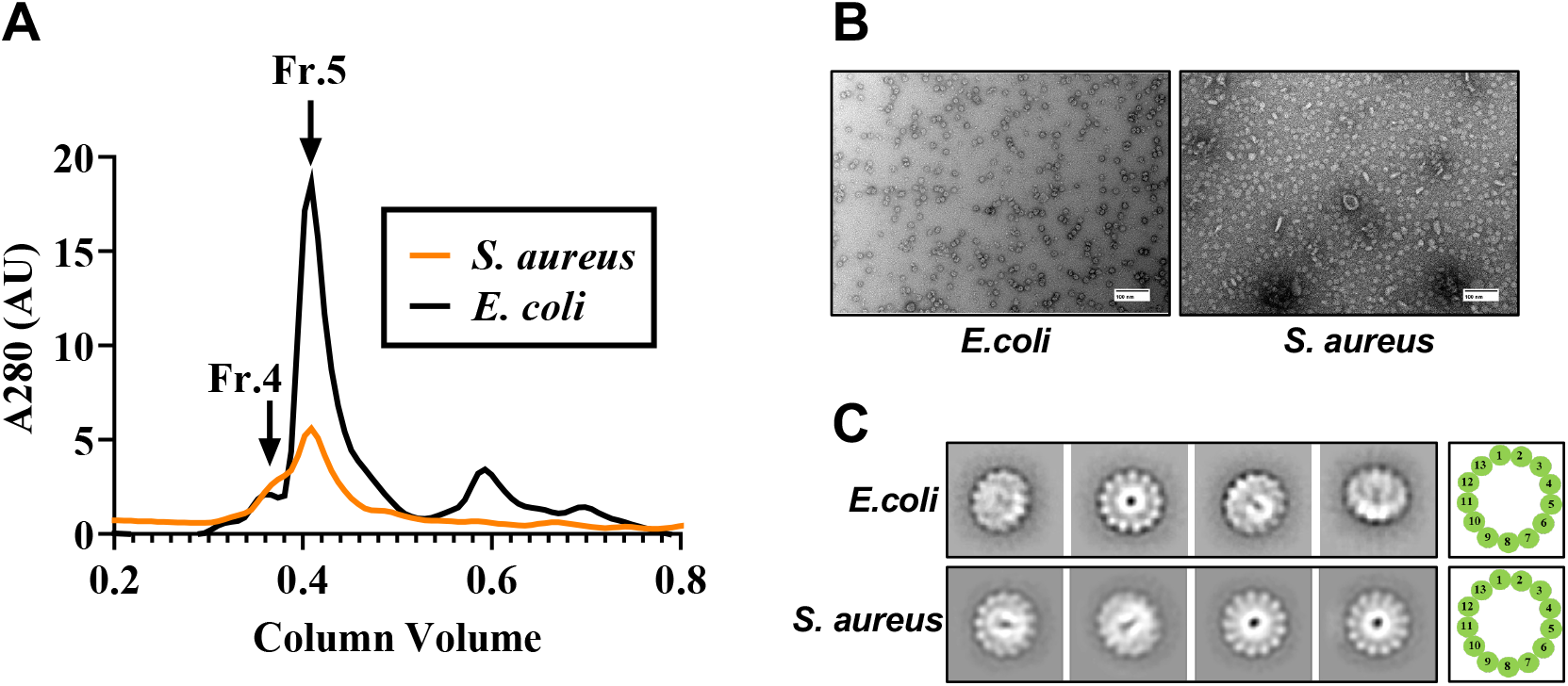
Expression of PP in *E. coli* and *S. aureus*. **(A)** SEC separation of wildtype PP expressed in *E. coli* (black) and *S. aureus* (orange). Fractions 4 and 5 are indicated. **(B)** Electron micrographs of negatively stained portals formed upon expression in *E. coli* and *S. aureus*. Scale bars = 100 nm. **(C)** 2D classification of negatively stained portals from *E. coli* and *S. aureus* expression.

### 2. Temperature dependence of portal formation

In our standard protocol for portal production from an *E. coli* lysate, the affinity purified protein was incubated at 37 °C for 3 h, as this was previously shown to promote oligomerization [26]. We hypothesized that the portal protein monomer adopts a quasi-stable conformation in solution that subsequently converts into an oligomer and that in the absence of other factors, elevated temperature (37°C) may promote the formation of tridecamer that represent the most stable structure in solution.

We tested this hypothesis expressing the portal protein in *E. coli* at 4 °C, 16 °C, 24 °C and 37 °C. The resulting portals were purified by Ni-NTA affinity and examined by negative stain EM (Supplementary Fig. S1), followed by 2D classification with cryoSPARC, which revealed multiple oligomeric states: octamers to tridecamers were observed at 4 °C and 16 °C, dodecamers to tridecamers at 25 °C, and 11-mers to tridecamers at 37 °C (Fig. 3A). The portals were further purified by SEC. Portals produced at 37 °C had a profile similar to that previously observed (Fig. 2A), whereas portals formed at other temperatures separated into multiple peaks (Fig. 3B). EM analysis and 2D classification of the different fractions revealed that the additional peak at fraction 4 contained portal doublets (Fig. 3C), whereas fraction 5 contained mostly separate tridecamers and some unclosed rings (Fig. 3C). Fraction 9 did not contain any visible portal rings, presumably constituting mainly monomeric protein. At 16 °C, an additional peak appeared at fraction 7; this peak also did not contain any ring-shaped structures, and was assumed to represent monomers or small oligomers. Overall, our SEC data show that monomers/small intermediates and doublets predominate at lower temperatures (Fig. 3D) while oligomers predominate at 37 °C.

**Figure 3.**
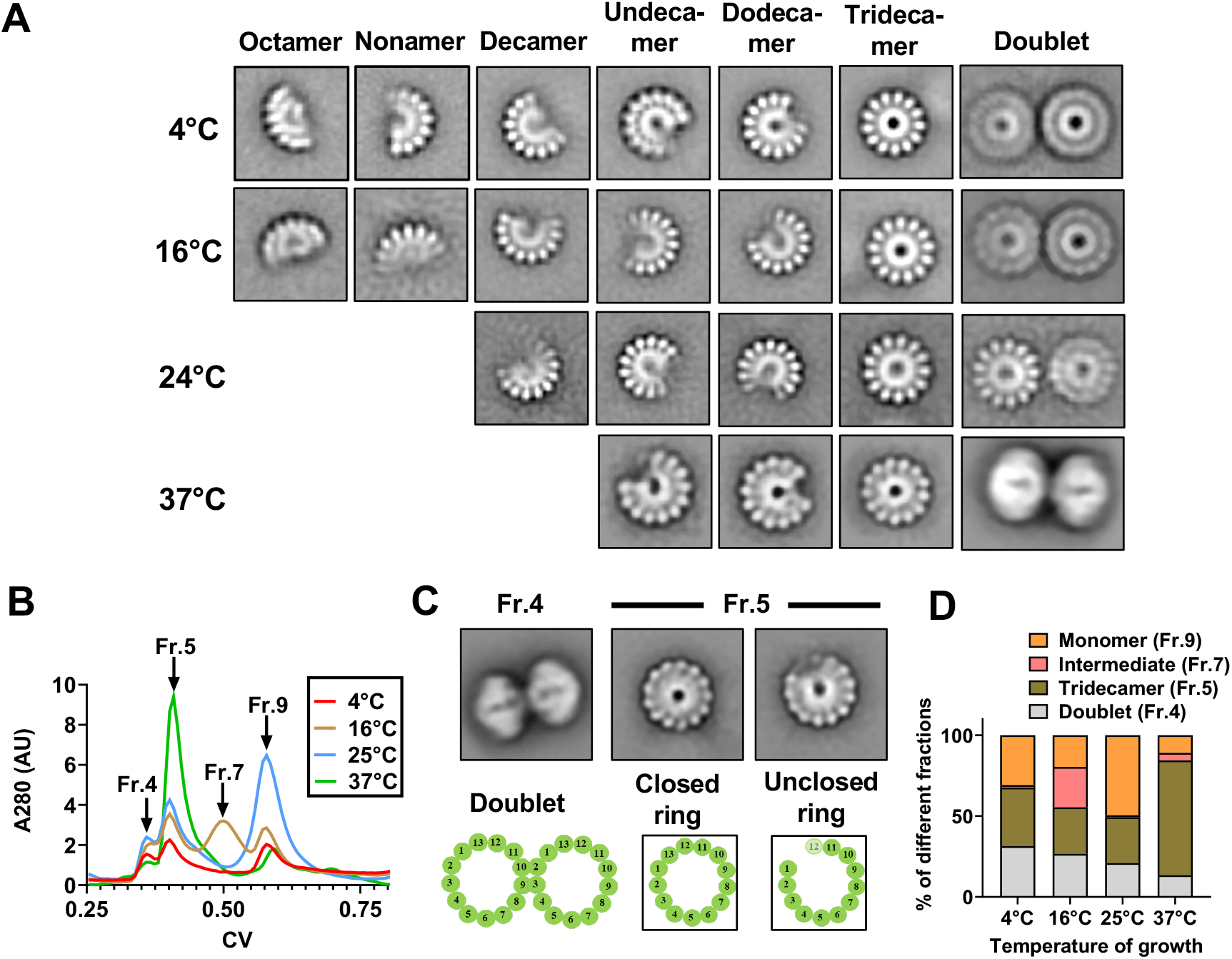
Portal formation at different temperatures. **(A)** 2D classification of negatively stained portals expressed at 4 °C, 16 °C, 25 °C or 37 °C, showing representative examples of closed and unclosed rings, as well as doublets. **(B)** SEC separation of portals purified from cells grown at 4 °C (red), 16 °C (orange), 25 °C (blue) or 37 °C (green). Peak fractions that were analyzed are indicated. **(C)** 2D classification of SEC fractions 4 and 5 from cells grown at 37 °C. **(D)** Histogram showing the ratio of SEC fractions 4 (doublet), 5 (oligomer), 7 (intermediate) and 9 (monomer) at different temperatures.

### 3. Assembly of portals with deletions of functional domains

We previously determined the structures of the isolated 80α portal tridecamers as well as the portal dodecamers existing in situ in capsids [26]. The PP monomer structure was similar in both cases, consisting of a wing domain (residues 1-257 and 358-431), an α-helical stem domain (258-285 and 336-357), a clip domain (286-335) and a C-terminal crown domain (432-511) (Fig. 4A). The N-terminus of PP were shown to interact with the CP [26], while the clip domain interacts with the connector (or adaptor) protein in the mature virion [10], and presumably with the large terminase (ORF41) during DNA packaging. The C-terminal crown domain forms a series of α-helices that protrude on the inside of the capsid, and was partially disordered in the 3D reconstructions of the portal [26]. An AlphaFold predicted model [29] of PP showed the extended C- and N-termini that were not resolved in cryo-EM map (Fig. 4B).

**Figure 4.**
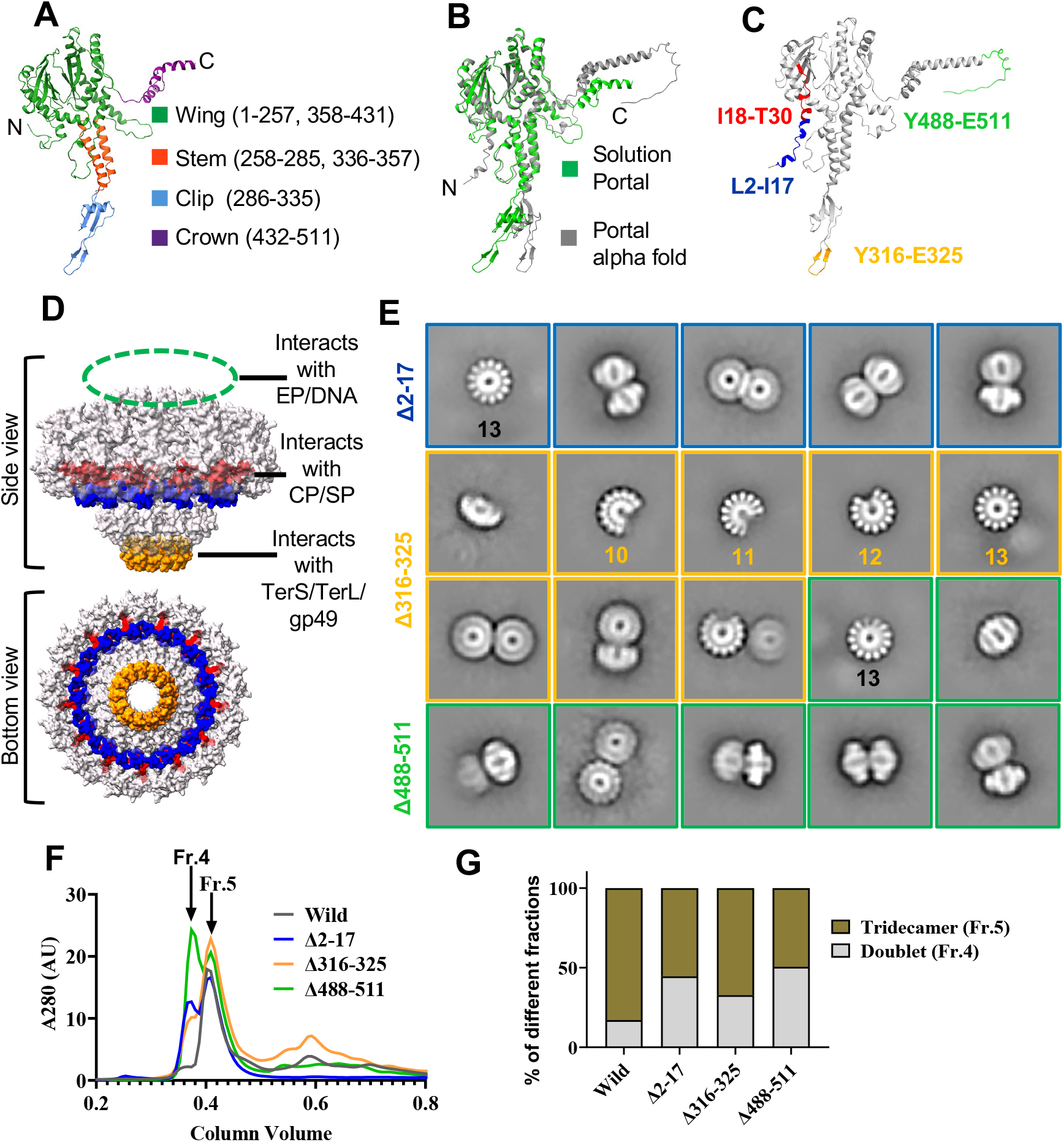
Analysis of portal deletions. **(A)** Ribbon diagram of 80α PP, determined by cryo-EM {Mukherjee et al., 2024, #232142}. The wing (green), stem (red), clip (blue) and crown (purple) domains are indicated. **(B)** Superposition of the PP oligomer determined by cryo-EM (green) on the AlphaFold model (gray). **(C)** AlphaFold model of the PP monomer, showing the sequences that were deleted in the deletion constructs PPΔ2-17 (L2-I17, blue), PP Δ2-30 (L2-T30, red), PPΔ316-325 (Y316-E325, yellow) and PPΔ488-511 (Y488-E511, green). **(D)** Deletions indicated on the PP oligomer (surface representation), colored as in (C), viewed from the side and from below. **(E)** 2D class averages of the three mutants PPΔ2-17, PPΔ316-325 and PPΔ488-511. The numbers indicate the number of PP subunits. **(F)** SEC separation of portals produced by PPΔ2-17 (blue), PPΔ316-325 (yellow) and PPΔ488-511 (green), compared to the wildtype (gray). Fractions 4 and 5 are indicated. **(G)** Histogram showing the ratio of fractions 4 (Doublet) and 5 (Tridecamer) in wild type and mutant portal proteins.

Based on this structure, we made PP expression constructs (Table 1) with the following deletions: PPΔ2-17 (N-terminus, wing domain), PPΔ2-30 (extended N-terminus, wing domain), PPΔ316-325 (clip domain) and PPΔ488-511 (crown domain) (Fig. 4C,D). The mutant portals were expressed in *E. coli*. PPΔ2-30 did not produce any portals, but the other three lysates were purified by Ni-NTA affinity, dialyzed and incubated at 37°C for 3–4 h to promote oligomerization as described earlier [26], followed by negative stain imaging (Supplementary Fig. S2) and 2D classification in cryoSPARC (Fig. 4E).

**Table 1.**
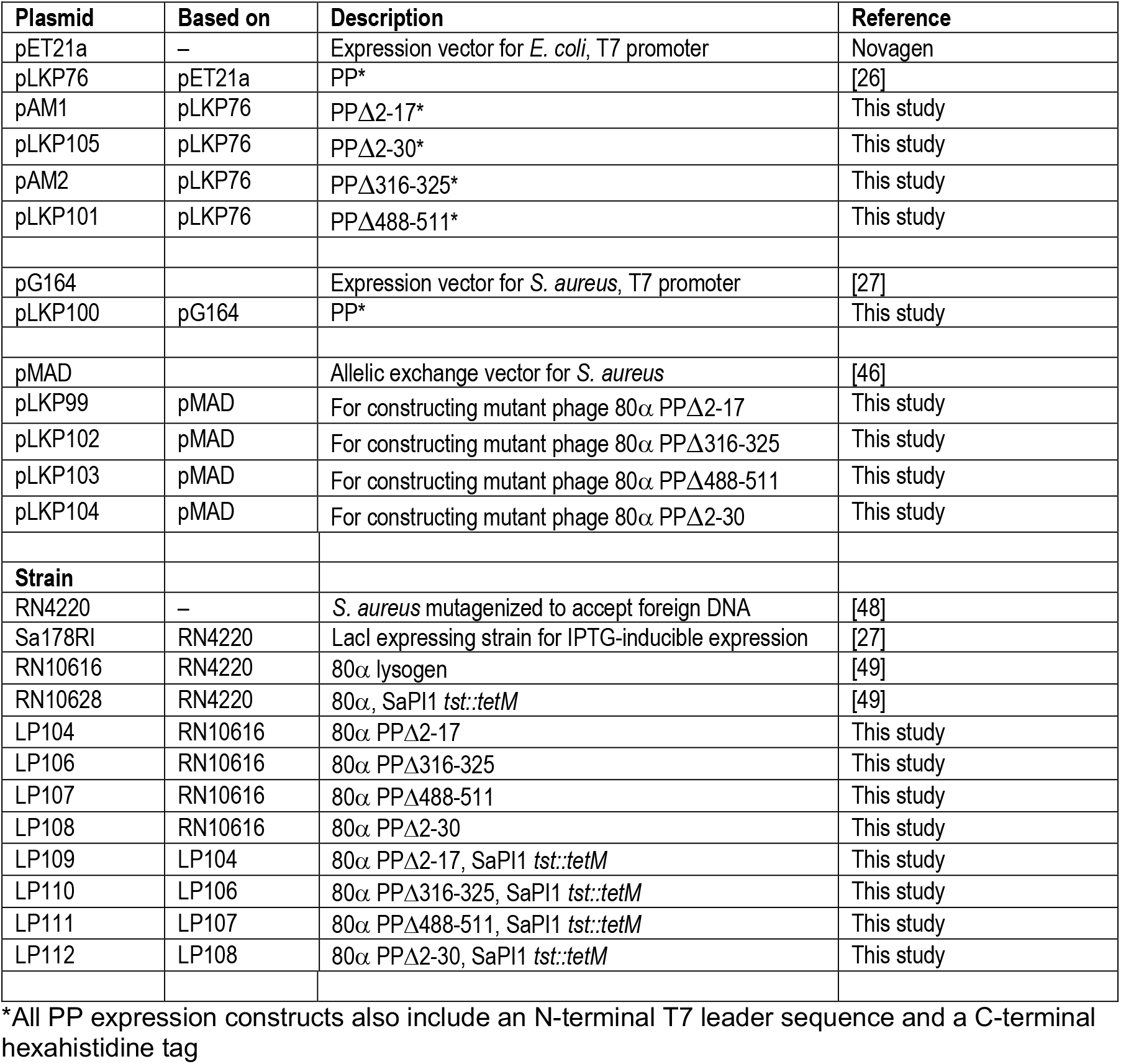
Plasmids and strains.

| Plasmid | Based on | Description | Reference |
| --- | --- | --- | --- |
| pET21a | – | Expression vector for *E. coli*, T7 promoter | Novagen |
| pLKP76 | pET21a | PP* | [26] |
| pAM1 | pLKP76 | PPΔ2-17* | This study |
| pLKP105 | pLKP76 | PPΔ2-30* | This study |
| pAM2 | pLKP76 | PPΔ316-325* | This study |
| pLKP101 | pLKP76 | PPΔ488-511* | This study |
| pG164 |  | Expression vector for *S. aureus*, T7 promoter | [27] |
| pLKP100 | pG164 | PP* | This study |
| pMAD |  | Allelic exchange vector for *S. aureus* | [46] |
| pLKP99 | pMAD | For constructing mutant phage 80α PPΔ2-17 | This study |
| pLKP102 | pMAD | For constructing mutant phage 80α PPΔ316-325 | This study |
| pLKP103 | pMAD | For constructing mutant phage 80α PPΔ488-511 | This study |
| pLKP104 | pMAD | For constructing mutant phage 80α PPΔ2-30 | This study |
| **Strain** |  |  |  |
| RN4220 | – | *S. aureus* mutagenized to accept foreign DNA | [48] |
| Sa178RI | RN4220 | LacI expressing strain for IPTG-inducible expression | [27] |
| RN10616 | RN4220 | 80α lysogen | [49] |
| RN10628 | RN4220 | 80α, SaPI1 *tst::tetM* | [49] |
| LP104 | RN10616 | 80α PPΔ2-17 | This study |
| LP106 | RN10616 | 80α PPΔ316-325 | This study |
| LP107 | RN10616 | 80α PPΔ488-511 | This study |
| LP108 | RN10616 | 80α PPΔ2-30 | This study |
| LP109 | LP104 | 80α PPΔ2-17, SaPI1 *tst::tetM* | This study |
| LP110 | LP106 | 80α PPΔ316-325, SaPI1 *tst::tetM* | This study |
| LP111 | LP107 | 80α PPΔ488-511, SaPI1 *tst::tetM* | This study |
| LP112 | LP108 | 80α PPΔ2-30, SaPI1 *tst::tetM* | This study |

\*All PP expression constructs also include an N-terminal T7 leader sequence and a C-terminal hexahistidine tag

PPΔ2-17 formed normal portals that were all tridecamers (Fig. 4E). PPΔ316-325 formed numerous unclosed rings, similar to those observed when the portal protein was expressed at 4 °C, while PPΔ488-511 produced normal looking tridecameric portals (Fig. 4E). By SEC, the mutant portals (PPΔ2-17, PPΔ316-325 and PPΔ488-411) all separated into two peaks at fractions 4 and 5, whereas the wildtype portals were only found in fraction 5 (Fig. 4F). Negative stain EM showed that fraction 4 in all cases contained portal doublets, while fraction 5 contained various closed and unclosed rings (Supplementary Fig. S2; Fig. 4G). PPΔ488-511 showed the highest propensity for doublet formation (Fig. 4F, 4G).

### 4. Stability of mutant portals

When the wildtype PP was analyzed by SDS-PAGE, a high molecular weight band appeared in addition to the 60 kDa monomer band when the samples were not boiled prior to loading on the gel (Fig. 5A), indicating the presence of an SDS-stable oligomer. The PPΔ2-17 and PPΔ488-511 mutant portals exhibited the same SDS-stable oligomer band, but the PPΔ316-325 mutant did not (Fig. 5A), indicating that this protein was inherently less stable.

**Figure 5.**
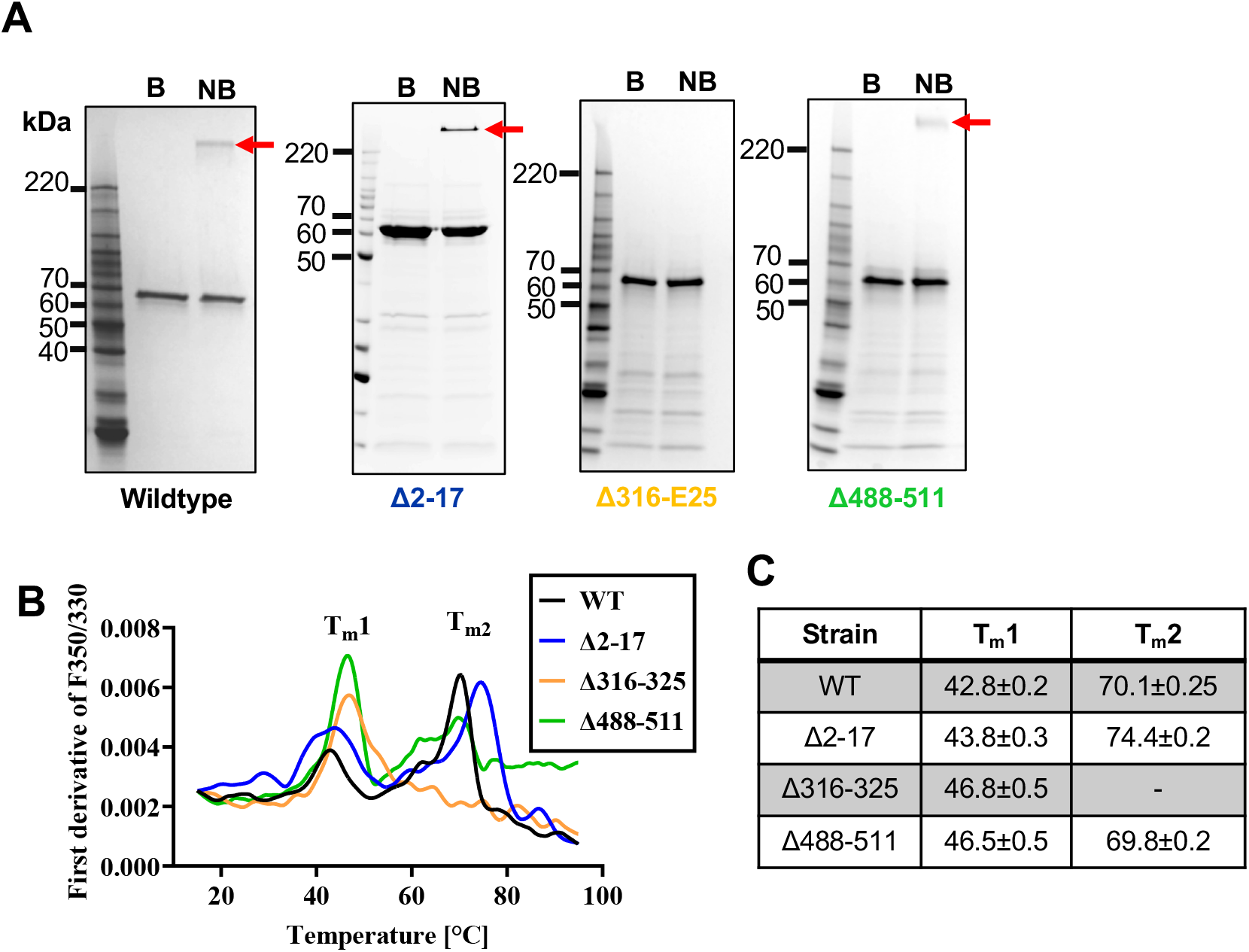
Thermal denaturation of PP. **(A)** SDS-PAGE of portals formed by wildtype PP and the PPΔ2-17, PPΔ316-325 and PPΔ488-511 deletion mutants, either boiled (B) or not boiled (NB) prior to separation. Bands corresponding to the PP oligomers are indicated by the red arrows. **(B)** Differential scanning fluorimetry of PP. The first derivative of the ratio of fluorescence emission at 350 nm and 330 nm (F350/F330) is plotted as a function of temperature for wildtype PP (black), PPΔ2-17 (blue), PPΔ316-325 (orange) and PPΔ488-511 (green). The two transition temperatures T_m_1 and T_m_2 are indicated. **(C)** Table of transition temperatures from (B). Mean and standard deviation of three measurements.

We further analyzed the thermal stability of each portal mutant using differential fluorimetry. Fraction 5 from wildtype PP and mutants PPΔ2-17, PPΔ316-325 and PPΔ488-411 were heated from 15 °C to 95 °C in 0.05 °C increments. The melting temperature (Tm) of each protein was determined using the ratio of tryptophan fluorescence emission at 350 nm and 330 nm as a measure of protein unfolding [30] and plotted as the first derivative of this ratio (Fig. 5B, 5C). This experiment showed that the wildtype portals exhibited transitions at 42.8 °C and 70.1 °C, consistent with an initial disruption of the oligomer at ≈43°C, followed by protein denaturation at ≈70°C (Fig. 5B, 5C). The PPΔ2-17 mutant had a similar biphasic profile, except that the upper peak was shifted up to 74.4 °C, indicating that the monomer of this protein is more stable than the wildtype. The PPΔ488-511 also displayed a biphasic profile, although in this case in the lower transition temperature was increased to 46.5 °C, indicating a slightly more stable oligomer formation, presumably due to the lack of the flexible C-terminal α-helices that may not contribute to the overall integrity of the oligomer. In contrast, PPΔ316-325 only had a single transition temperature at 46.8 °C (Fig. 5B, 5C), suggesting that this mutant was highly unstable and denatured at the same temperature as the oligomer falls apart.

### 5. Structure of the PPΔ316-325 mutant

The portals formed by the PPΔ316-325 mutant were purified by SEC and subjected to cryo-EM (Fig. 6A) and 3D reconstruction with cryoSPARC. 2D classification revealed several classes that looked like unclosed rings with various numbers of subunits (Fig. 6B). The particles were subjected to ab initio model generation and 3D classification with 20 classes, yielding various types of closed and unclosed rings with from 8 to 14 subunits (Fig. 6C). Some of the unclosed rings displayed a distinct twist rather than the typical planar rings. The most common closed ring was a tridecamer, but a tetradecameric class was also observed, suggesting that the unclosed rings formed by the mutant can be stabilized through incorporation of additional subunits in solution.

**Figure 6.**
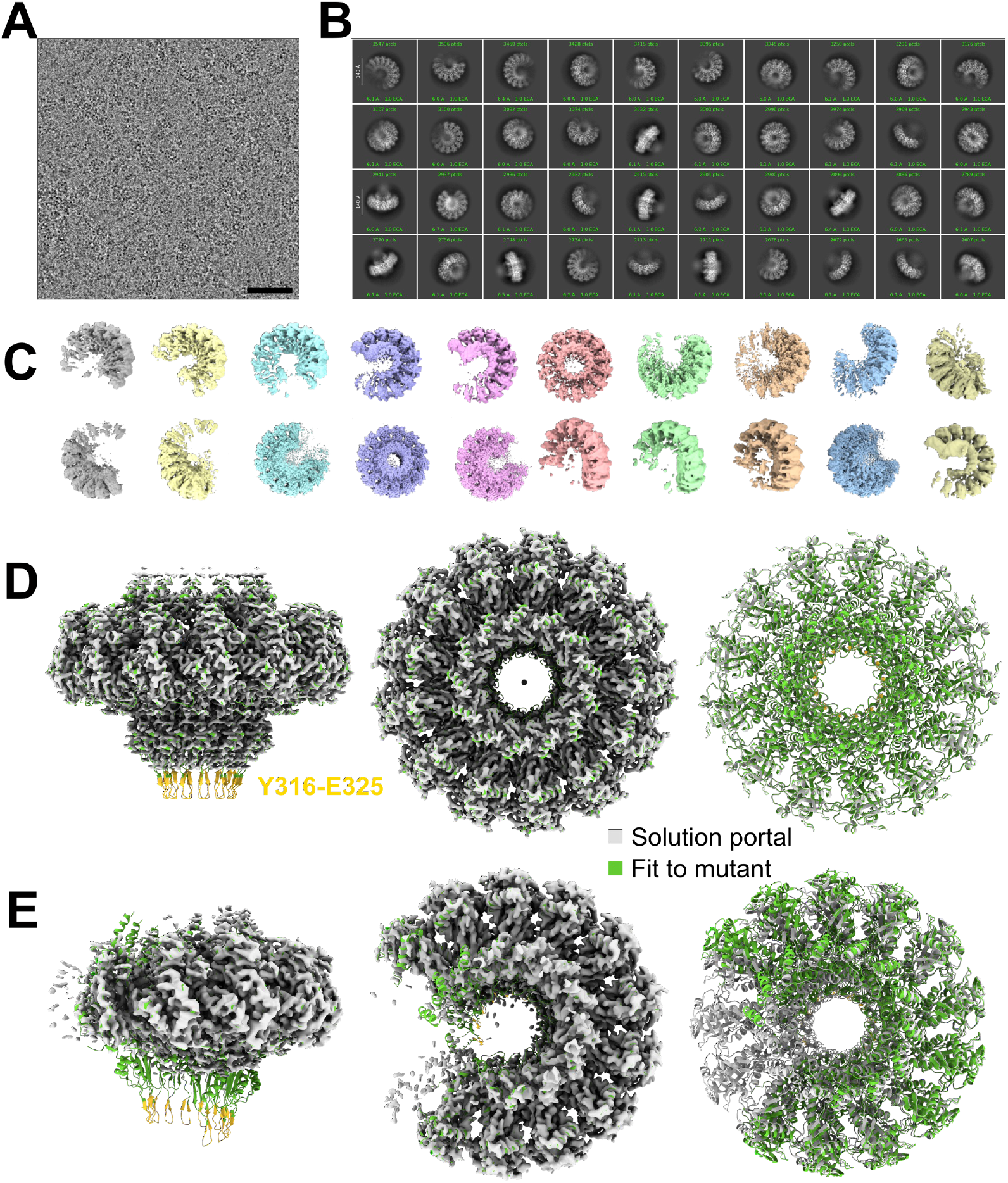
3D reconstruction of portals from PPΔ316-325 mutant. **(A)** Cryo-electron micrograph of PPΔ316-325 portals. Scale bar = 50 nm. **(B)** Examples of 2D classes from cryo-EM of PPΔ316-325 portals. **(C)** 3D classes from heterogeneous refinement of large oligomers, showing the range of variability in portal structures (default ChimeraX coloring, 5σ threshold). **(D)** The tridecamer reconstruction (4σ threshold) and **(E)** the unclosed undecamer reconstruction (5σ threshold) with the previously determined wildtype PP monomer atomic model fitted individually for each subunit (green ribbon diagram with residues Y316-E325 in yellow) or fitted as a complete tridecamer (gray ribbon diagram).

The classes were then subjected to heterogeneous refinement. The class representing the closed tridecamer reached a resolution of 2.9 Å (Fig 6D, Supplementary Fig. S3). The previously determined model of the portal [26] was fitted into this map, showing that the structures were essentially identical, except for the missing clip domain (residues 316-325)(Fig. 6D). Another class representing an incomplete portal ring containing 11 subunits, apparently missing two subunits of a tridecamer, reached 3.4 Å resolution (Fig. 6E). When the previous atomic model was fitted into this map, it was clear that the map was not only missing the clip domain residues that had been deleted, but were also missing large portions of the stem domain (residues 258-357), indicating disorder or flexibility in this region (Fig. 6E). The other classes could not be refined to high resolution, due to a combination of heterogeneity and strong orientation bias.

### 6. Formation of virions in phage with portal deletions

The same four mutations (Δ2-17, Δ2-30, Δ316-325 and Δ488-511) were introduced into the ORF42 gene in the 80α lysogen strain RN10616, generating phages 80αPPΔ2-17 (strain LP104), 80αPPΔ2-30 (LP108), 80αPPΔ316-325 (LP106) and 80αPPΔ488-511 (LP107) (Table 1). We also introduced SaPI1 *tst::tetM* into the same strains by transduction, generating strains LP109 (80αPPΔ2-17), LP112 (80αPPΔ2-30), LP110 (80αPPΔ316-325) and LP111 (80αPPΔ488-511) (Table 1).

The mutant strains were induced with mitomycin C as previously described, and the resulting crude lysates were examined by negative stain EM (Fig. 7A). Mutant 80αPPΔ2-17 formed normal virions with filled capsids in the absence of SaPI1 (strain LP104), and normal transducing particles with small capsids in the presence of SaPI1 (LP109), showing that residues 2-17 are not required for capsid assembly, DNA packaging and tail attachment (Fig. 7A). In contrast, mutant 80α PPΔ2-30 formed few capsids-like structures and no virions both without (LP108) or with (LP112) SaPI1, consistent with the lack of portal formation described above (Fig. 7A). Mutant 80α PPΔ316-325 (strain LP106) formed only procapsids and did not attach tails, consistent with a defect in interacting with terminase and with the connector complex required for tail attachment (Fig. 7A). In the presence of SaPI1 (strain LP110), these procapsids were predominantly small (Fig. 7A). Mutant 80α PPΔ488-511 produced normal looking virions with filled capsids, both without (strain LP107) and with (LP111) SaPI1 (Fig. 7A). (In strain LP111 the capsids were small.)

**Figure 7.**
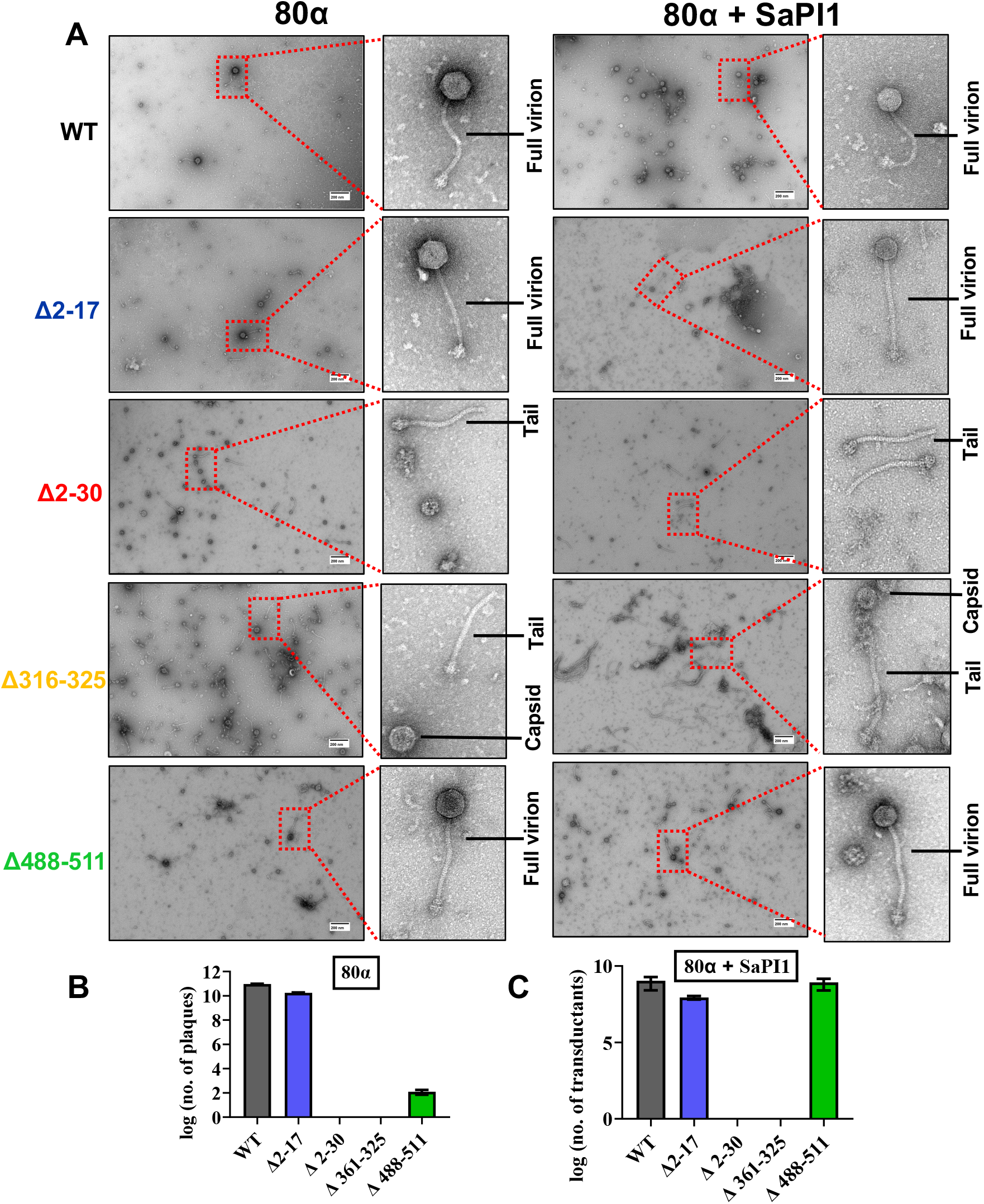
Electron microscopy of mutant phage. **(A)** Negatively stained material produced by the WT and mutant 80α lysogens 80αPPΔ2-17 (LP104), 80αPPΔ2-30 (LP108), 80αPPΔ316-325 (LP106) and 80αPPΔ488-511 (LP107). and the corresponding SaPI1*tst:tetM*-containing lysogens LP109 (80αPPΔ2-17), LP112 (80αPPΔ2-30), LP110 (80αPPΔ316-325) and LP111 (80αPPΔ488-511). Scale bars = 200 nm. The panels to the right show magnified views with pertinent features indicated. **(B)** Phage titers, plotted as log_10_ of the plaque forming units (pfu) of wildtype (WT) and mutant phage lysates. **(C)** Transducing titers for SaPI1 particles produced by wildtype and mutant phages, expressed as log_10_ of the number of tetracycline-resistant transductants (colony forming units).

We also examined the phage and transducing titers of the mutant strains. Consistent with the observations above, 80αPPΔ2-17 had near-wildtype phage titers (a tenfold reduction) (Fig. 7B), and a fivefold reduction of SaPI1 transducing titer (Fig. 7C). Mutants 80αPPΔ2-30 and 80αPPΔ316-325 had zero phage and transducing titers, consistent with the observed defects in capsid assembly and terminase recruitment, respectively. However, mutant 80αPPΔ488-511 had a distinct phenotype: although the virions looked normal (Fig. 7A), the phage titer was reduced 10^8^-fold compared to the wildtype (Fig. 7B). However, in the presence of SaPI1, the transducing titer was near-identical to the wildtype (Fig. 7C). This phenotype is consistent with a reduced burst size that is insufficient to form plaques, whereas SaPI1 transducing titers, which are not dependent on replication, are unaffected.

We hypothesized that the observed defect might be due to altered interaction of the C-terminal crown domain of PP with the genome in the 80α Δ488-511 mutant, leading to either an unstable DNA that is prone to premature ejection or a defect in ejection itself. This was tested using a DNA ejection assay that we previously described [31]. Purified virions were heated to 68°C for up to 5 min, which was previously shown to induce DNA ejection. The ejected DNA is degraded with DNAse I. After inactivating the DNase with EDTA, the remaining capsids are treated with proteinase K to disrupt the capsids, followed by quantitation of the remaining DNA by agarose gel electrophoresis.

This experiment showed that there was a slightly increased tendency of DNA ejection in the 80α PPΔ488-511 mutant virions produced both in the absence and presence of SaPI1 after 2 min at 65 °C (Fig. 8A, 8B). At 5 min, the difference was less pronounced, since most of the DNA was already ejected at this time point (Fig. 8A, 8B). However, the slightly increased ejection at 2 min was insufficient to explain the big difference in phage titer (Fig. 7B).

**Figure 8.**
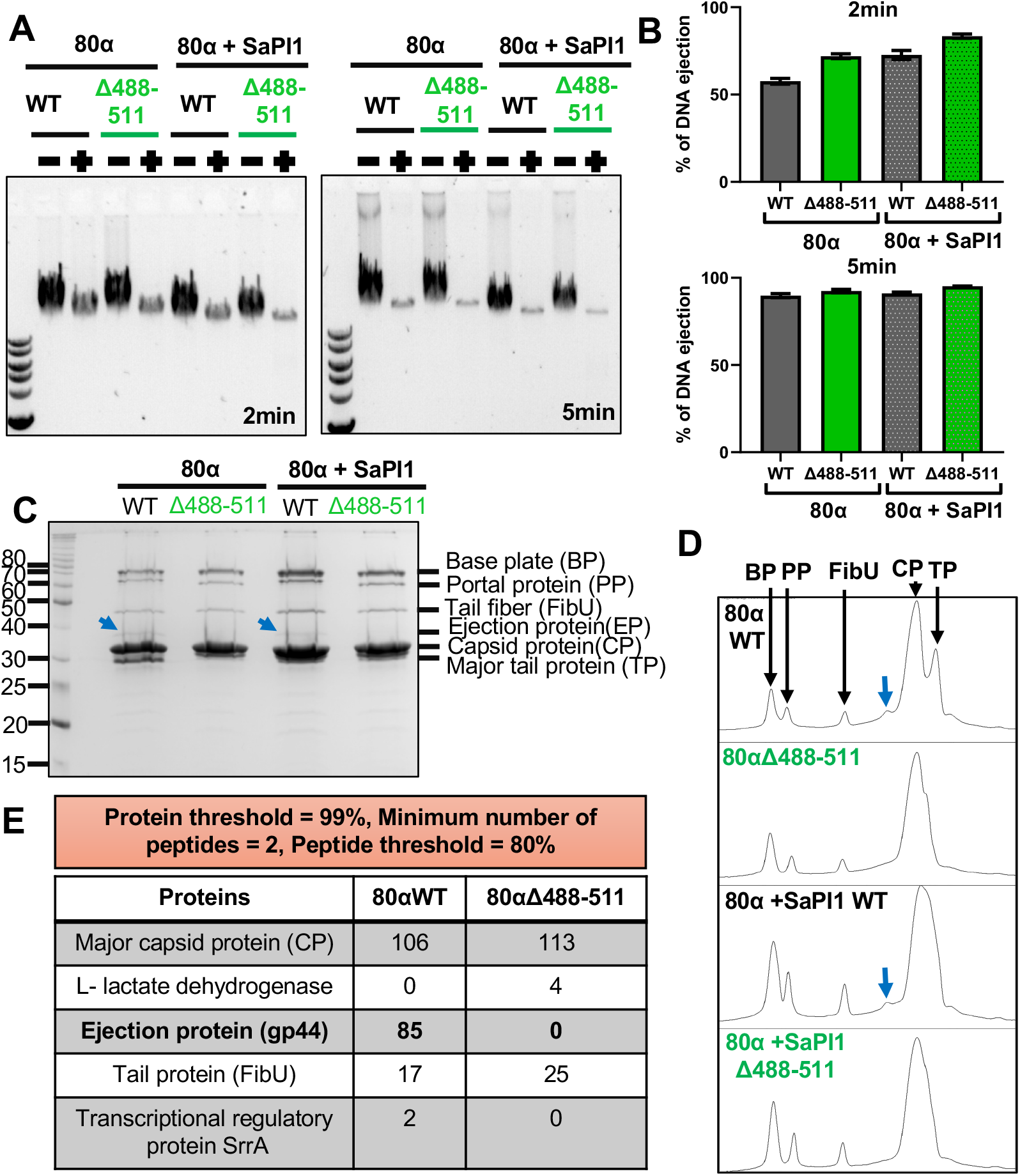
DNA ejection and protein composition of PPΔ488-511 mutant phage. **(A)** Agarose gel of DNA ejected from 80α WT and PPΔ488-511 virions and SaPl1 particles after 2 min (left) or 5 min (right) incubation at 68 °C (+) and 4 °C (–). **(B)** Quantitation of DNA ejection in 80α virions and SaPl1 particles after 2 or 5 min incubation at 68 °C, measured by densitometry of the DNA gels and expressed as the percentage DNA ejected relative to samples kept at 4 °C. **(C)** SDS-PAGE of wildtype and PPΔ488-511 phage and SaPl1 particles. Bands corresponding to baseplate proteins (BP), PP, upper tail fiber (FibU), CP and major tail protein (TP) [47] are indicated. A faint band corresponding to EP (gp44) can be seen in the wildtype 80α and SaPl1 particles [31]. (D) Densitometric plot of the gel in (C) measured by lmageJ. Position of EP is indicated by the arrow. **(E)** Table of proteins identified by MS in the EP band from (C) in wildtype 80α and 80α PPΔ488-511, listing the number of detected peptides.

### 7. Presence of the ejection protein gp44

Previous results had shown that minor capsid protein gp44 is an ejection protein (EP) that protects the phage DNA post injection [16]. While phage titers are greatly reduced in an 80αΔ44 mutant, SaPI1 titers are unaffected, similar to the phenotype described above for the 80αPPΔ488-511 mutation [31].

We therefore asked whether the observed phenotype for the 80αPPΔ488-511 mutant was due to the presence or absence of EP. When the phage and SaPI1 particles formed by this mutant were run on a 15% SDS-PAGE gel, there was a band consistent with EP present (at apparent molecular weight of 39kDa) in the wildtype that was absent from the PPΔ488-511 mutant (Fig. 8C, D). The corresponding segments were excised from the gel and subjected to LC-MS analysis after trypsin digestion, which confirmed that the band observed in the wildtype contained EP, while the 80αPPΔ488-511 mutant lacked this protein (Fig. 8E). This result suggests that the crown domain of PP interacts with gp44 and is required for incorporation of EP into phage and SaPI1 particles.

## DISCUSSION

Although portals found in phage capsids are invariably dodecamers [13, 14], portals produced after overexpression in *E. coli* frequently adopt other oligomeric states, including the tridecamer we observe for the 80α PP [26]. Tridecamers were also observed for *Bacillus* phages SPP1 and ϕ29 [32–34]. *E. coli* phage T7 formed a mixture of tri- and dodecamers [35, 36], while *Salmonella* phage P22 formed undecamers and dodecamers [34]. A distribution from 11-to 14-mer was observed for herpes simplex virus 1 [37]. In contrast, the portals of *E. coli* phages P2 (gpQ) [38], lambda (gpB) [39] and T3 (gp8) [40] formed dodecamers even upon overexpression, potentially implicating host-specific factors in correct oligomerization. Why do portals form these apparently unproductive oligomeric states, and what leads to the formation of the closed dodecamer? Are the dodecameric portals initially formed as tridecamers or are there additional factors in the native system in vivo that shift the assembly towards dodecamers before the unproductive tridecamers are formed?

In this study, we have examined this phenomenon further. We show that portal formation is temperature sensitive (Fig 3), but does not appear to be dependent on *S. aureus*-specific factors, as we observed no difference in portal structure when protein was expressed in *S. aureus* compared to *E. coli* (Fig. 2). Our data suggest that portals are initially formed as incomplete, unclosed rings with various numbers of subunits. We did not observe oligomers smaller than octamers, either because these are transient and only present in very low numbers, or because they were too small or heterogeneous to be classified in cryoSPARC. At low temperatures, these intermediates are kinetically trapped, while increasing temperature promotes their conversion into closed tridecamers. Additionally, at lower temperatures, rings have a tendency to dimerize, probably due to interactions between incomplete rings (Fig. 9).

**Figure 9.**
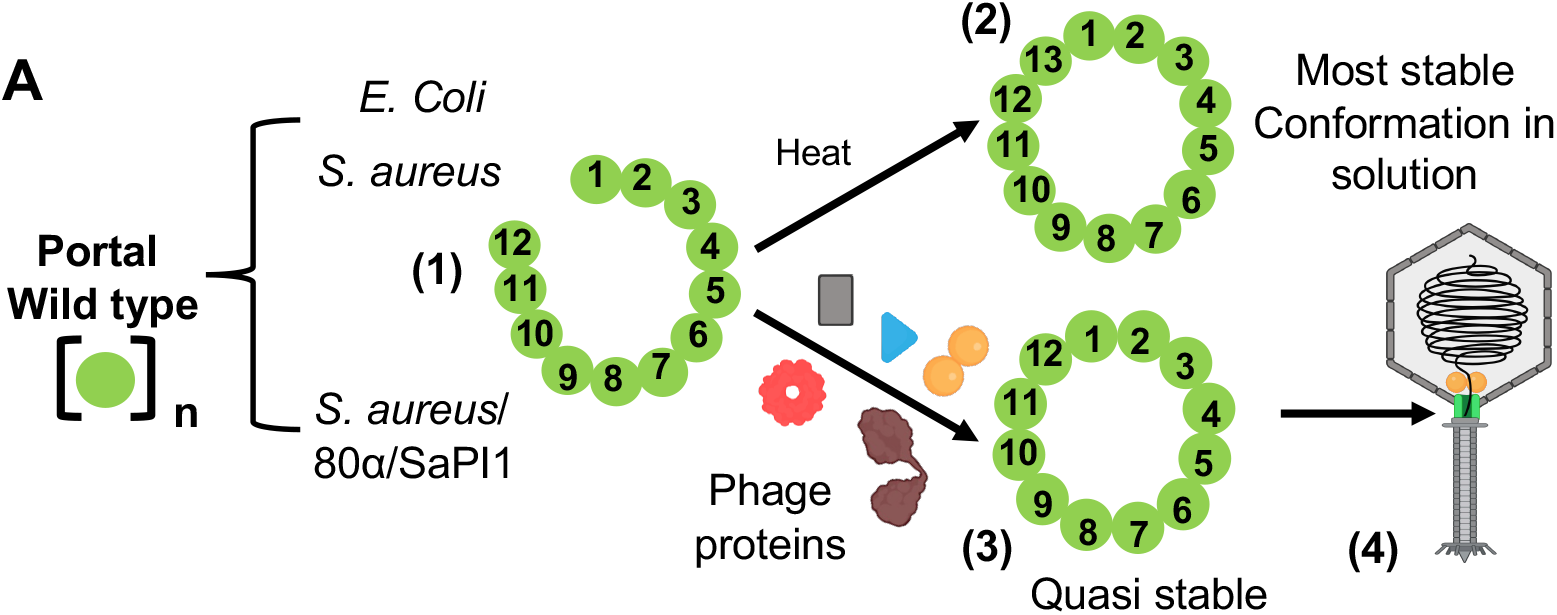
Model of portal assembly and incorporation. Portal ring formation in the presence and absence of other 80α phage proteins. (1) Formation of unclosed rings with a variable number of subunits; (2) Heat converts unclosed rings to tridecamers, the most stable state in solution; (3) The presence of other phage proteins converts unclosed rings to quasi-stable dodecamers; (4) Only dodecamers are incorporated into capsids capable of packaging DNA.

We predict that during a normal infection, in the presence of other phage proteins, these unclosed rings are converted into dodecamers (Fig. 9). The dodecameric portal may represent a metastable state, which could be important for its conformational plasticity during capsid assembly, DNA packaging, and DNA ejection. The SPs are most likely involved in this process: an interaction between PP and SP is expected by comparison with other systems [41, 42], and in P22, PP oligomerization *in vivo* was shown to be driven by SP [43, 44], although it is not clear whether this interaction affected the oligomeric state per se. Our data also indicate that closure of the ring is dependent on the clip domain that is expected to interact with the terminase during DNA packaging and with the connector during head completion and subsequent tail joining, potentially implicating the terminase in portal formation and incorporation.

Interestingly, portals lacking C-terminal residues 488-511 did not incorporate EP (gp44)(Fig. 8C), which we previously showed to protect the ends of the phage genome post injection [16, 31]. EP is not absolutely essential for viability of 80α and is dispensable for SaPI1 transduction, but increases phage burst size. This explains the observed difference in 80α phage and SaPI1 transducing titers with the PPΔ488-511 mutant. The lack of EP in this mutant suggests that this protein is incorporated into capsids by interaction with the portal. No defect in portal formation or assembly was observed in this mutant, however.

We previously showed that the flexible N-terminal domain of PP interacts with the capsid, and it was suggested to play an important role in portal incorporation and capsid assembly [26, 45]. Surprisingly, deletion of PP residues 2-17 had little effect on assembly and viability. In contrast, the longer deletion PPΔ2-30 failed to form portals, and the 80αPPΔ2-30 strain produced no capsids (Fig. 7A). This was somewhat surprising, since we know that capsids can be formed in the absence of portals. Perhaps the misfolded mutant PP prevented assembly by capturing other structural proteins.

Based on the above results, we can propose the following sequence of events leading to the production of functional procapsids (Fig. 9): (1) Assembly of loosely associated PP oligomers and unclosed rings; (2) In the absence of other phage proteins, the end point is a non-functional tridecamer, representing the most stable conformation in solution; (3) Interaction with other phage proteins, leads to the formation of a closed dodecamer. EP might be incorporated at this stage as well, but is not essential; (4) Capsid assembly and incorporation of dodecameric portals into capsids.

The incorporation of portals is a key step in the formation of functional procapsids that are capable of packaging the genome and interact with the DNA. This process is highly efficient in vivo, but has proven to be difficult to reproduce in vitro, highlighting the complex dynamics and finetuned interplay between the various phage proteins that have evolved over eons.

## MATERIALS AND METHODS

### 1. Cloning in *E. coli* and *S. aureus*

The *E. coli* plasmid pLKP76, described previously [26], encodes the 80α PP (gp42) with a N-terminal T7 leader tag and a C-terminal hexahistidine tag under the T7 promoter (Table 1). This plasmid was used as a template for deletion mutagenesis using the In-Fusion Cloning Kit (Takara). The resulting plasmids pAM1, pLKP105, pAM2 and pLKP101 express PP deletions PPΔ2–17, PPΔ2–30, PPΔ316–325 and PPΔ488–511, respectively (Table 1). All constructs include an N-terminal T7 leader tag and a C-terminal hexahistidine tag.

For cloning in *S. aureus*, the pG164 shuttle vector was used [27]. The portal protein gene (ORF42) was amplified from the pLKP76 vector and subcloned into pG164 using the In-Fusion Cloning Kit (Takara). The resulting plasmid pLKP100 was introduced into Sa178RI competent cells by electroporation, followed by selection with chloramphenicol.

### 2. Expression and purification of portals

The PP-containing plasmids were expressed in *E. coli* BL21 (DE3) as described previously [26] and the PP was purified by affinity on Ni-NTA agarose resin (Invitrogen) followed by dialysis overnight at 4 °C in 25 mM HEPES buffer pH 7.4, 1 M NaCl. For expression in *S. aureus*, cells were grown at 32 °C in Tryptic Soy Broth (TSB) until the optical density at 600 nm (OD_600_) reached 0.5, followed by induction with 0.5 mM IPTG. After growing for another 3 h, the cells were harvested by centrifugation, resuspended in 25 mM HEPES pH 7.4, 1 M NaCl and lysed using an Avestin Emulsiflex high pressure disruptor. The protein was purified using Ni-NTA agarose resin (Invitrogen) as above.

For expression of PP at different temperatures, *E. coli* BL21 (DE3) cells harboring pLKP76 was cultured in LB broth at 37°C until the optical density at 600 nm (OD_600_) reached 0.5. Protein expression was induced by the addition of 1mM IPTG, after which the cultures were immediately transferred to incubators set at different temperatures. The cells were further incubated at 25 °C for 6 h, or at 16 °C and 4 °C for 16 h. Cells were harvested, and the recombinant protein was purified using Ni–NTA agarose resin (Invitrogen) at 4 °C.

Purified proteins were subsequently applied to a Superdex™ 300 Increase size-exclusion chromatography column (GE Healthcare). Eluted fractions were collected and analyzed by SDS– PAGE.

### 3. Differential scanning fluorimetry

Temperature-dependent unfolding of wild-type and mutant portal proteins was analyzed using a Prometheus NT48 nanoDSF instrument (NanoTemper Technologies). Proteins were dialyzed against 25 mM HEPES buffer (pH 7.4) containing 0.1 M NaCl. Tryptophan fluorescence emission was recorded over a temperature range of 15–95°C with an increment of 0.05°C. The thermal unfolding profile was plotted as the first derivative of the ratio of fluorescence emission intensities at 350 nm and 330 nm (F350/F330) versus temperature.

### 4. Genetic manipulation in bacteriophage

Different deletion mutants of the ORF42 gene were subcloned into the pMAD allelic exchange vector [46] using the In-Fusion Cloning Kit (Takara). The recombinant plasmids were transformed into *E. coli* Stellar cells (Takara) and verified by DNA sequencing. Transformation and allelic exchange of the pMAD vectors carrying various portal protein mutants were performed in the *S. aureus* 80α lysogen strain RN10616 as described previously [47]. Successful recombinants were confirmed by sequencing. The resulting 80α mutant lysogens 80α PPΔ2-17, 80α PPΔ316–325, 80α PPΔ488–511, and 80α PPΔ2–30 were designated LP104, LP106, LP107, and LP108, respectively (Table 1). The 80α portal mutant strains were subsequently transduced with SaPI1 *tst::tetM* particles produced by 80α lysogen strain RN10628 to generate SaPI1-containing 80α derivatives with the corresponding portal protein deletions. The resulting strains, designated LP109, LP110, LP111, and LP112, carried 80α mutant lysogens 80α PPΔ2-17, 80α PPΔ316– 325, 80α PPΔ488–511, and 80α PPΔ2–30, respectively (Table 1).

### 5. Purification of bacteriophage, phage titer and transduction assay

80α virions and SaPI1 transducing particles were purified by PEG precipitation and CsCl gradient centrifugation as described previously [8, 26]. The phage titers of the portal deletion mutants were determined by plating on *S. aureus* strain RN4220 and compared to that of the wild-type 80α phage. Similarly, the transduction titers of SaPI1 particles produced in 80α lysogens carrying the portal protein deletions were measured by counting tetracycline-resistant colonies after mixing the SaPI1 particles with *S. aureus* strain RN4220, and compared to those of the wild-type SaPI1 particles.

### 6. In vitro DNA ejection assay

Purified 80α virions and SaPI1 particles were incubated at 68 °C for 2 or 5 min in a thermocycler to release encapsidated DNA. The released DNA was then treated with 40 µg/ml DNase I for 1 h at 37°C, followed by enzyme inactivation with 10 mM EDTA (pH 8.0) and incubation at room temperature for 30 min. Subsequently, 1 mg/ml Proteinase K was added, and the samples were incubated at 55 °C for 1 h in a thermocycler to ensure complete release of capsid-protected phage DNA. The resulting DNA samples were analyzed by electrophoresis on a 0.7% agarose gel run at 70 V for 2 h. The percentage of DNA released, relative to the unheated control sample, was quantified by densitometric analysis using ImageJ.

### 7. Mass spectrometry

Purified 80α phage particles were boiled in Laemmli sample buffer and separated by 15% SDS– PAGE, followed by staining with colloidal Coomassie Blue. The band corresponding to the expected mass of gp44 (38.5 kDa) was excised from the gel and subjected to trypsin digestion. The resulting peptide mixture was analyzed by liquid-chromatography mass spectrometry (LC-MS) using a Q Exactive HF-X Orbitrap spectrometer (Thermo Scientific) and analyzed by Scaffold 5.

### 8. Electron microscopy

Purified portal, 80α virion and SaPI1 particle samples were applied to glow-discharged grids with ultrathin carbon layered on lacey carbon (Electron Microscopy Sciences), negatively stained with 1% uranyl acetate and examined using either an FEI F20 transmission electron microscope (TEM) with a Gatan OneView detector, operated at 200 kV, or a JEOL 1400 HC Flash TEM with an AMT NanoSprint detector, operated at 120 kV. The micrographs of purified portal proteins were subjected to 2D classification using CryoSPARC v4.6.0.

For cryo-EM analysis, the PP oligomers were separated at 154,000 g for 2 h on a 5-20% sucrose gradient in dialysis buffer. Fractions containing the highest concentration of portal protein oligomers were dialyzed overnight in 25mM HEPES, pH-7.4, 0.1M NaCl and concentrated using an Amicon Ultra 100 kDa cut-off membrane (Millipore). Cryo-EM samples were prepared on glow-discharged copper C-flat R2/1 grids using a Vitrobot Mark IV, and imaged in a Thermo Fisher Glacios 2 microscope equipped with a Falcon 4i detector, operated at 200 kV. A total of 17,157 38-frame movies were collected at a magnification of 190,000x, corresponding to a pixel size of 0.7159 Å, and a total electron dose of 40 e^−^/Å^2^. (Table S1).

### 9. Reconstruction of portals

Cryo-EM images were processed in CryoSPARC v4.7.1, including preliminary Live processing during collection. Initial particle picking used a ring template with 50 Å internal and 180 Å external diameters. 2D classes were subsequently used for template-based picking of 6.4 million particles. Further 2D classification, *ab initio* 3D reconstruction, and heterogeneous refinement yielded a single 3.37 Å-resolution undecamer portal reconstruction. Picking was repeated on denoised micrographs using templates generated from the undecamer reconstruction and from a tetramer created from the solution structure atomic model (PDB: 8V8B) using ChimeraX v1.8. After curating the two sets of particles, the remaining 1.7 million particles in total were combined, reconstructed into 20 *ab initio* classes, and heterogeneously refined. The resulting classes represented various oligomers as small as 2-3 subunits. The six classes corresponding to larger oligomers of six or more subunits (951,063 particles) were subjected to further heterogeneous refinement into 20 classes, followed by homogeneous refinement. The single tridecamer class (62,289 particles) and the best undecamer class (64,969 particles) were further refined including CTF and aberration correction, applying either C13 symmetry averaging or relaxation. The C13-symmetrical tridecamer reached a final resolution of 2.88 Å, and the asymmetric undecamer class reached 3.39 Å. The volumes were colored according to local resolution estimates from CryoSPARC using ChimeraX v1.10.

Subsequent analysis of the maps and atomic models was done with ChimeraX v1.10. Because of the similarity between the tridecamer reconstruction and the previous solution structure, the PP monomer (PDB: 8V8B) was rigid-body fitted to each subunit of the tridecamer reconstruction for comparison. The root-mean-square deviation between the WT solution structure and the resulting tridecameric mutant structure was 0.38 Å. Similarly, eleven copies of the WT monomer structure were fitted into the undecamer mutant reconstruction.

## ACKNOWLEDGEMENTS

Electron microscopy of negatively stained samples was done at the UAB High Resolution Imaging Facility (HRIF) and at The University of Alabama, Tuscaloosa (UA) Core Analytical Facility. We appreciate the assistance of Melissa Chimento at HRIF and Robert Holler at UA with the electron microscopy. Cryo-EM was done in the UAB Cryo-EM Facility (CEMF), supported by National Institutes of Health (NIH) grant S10 OD024978 to T.D. We are grateful for the assistance of Dr. Kyoko Kojima in the UAB Mass Spectrometry/Proteomics (MSP) Shared Resource for assistance with the MS experiments, and Wendy Yang in the UAB X-ray Crystallography Shared Facility for help with Differential Scanning Fluorimetry. HRIF, MSP and CEMF were supported by the O’Neal Comprehensive Cancer Center at UAB (NIH grant P30 CA013148) and the UAB Institutional Research Core Program (IRCP). This work was supported by NIH grant R01 AI083255 to T.D.

