## Supplementary Figures for "Formation and role of the portal of *Staphylococcus aureus* bacteriophage 80α"

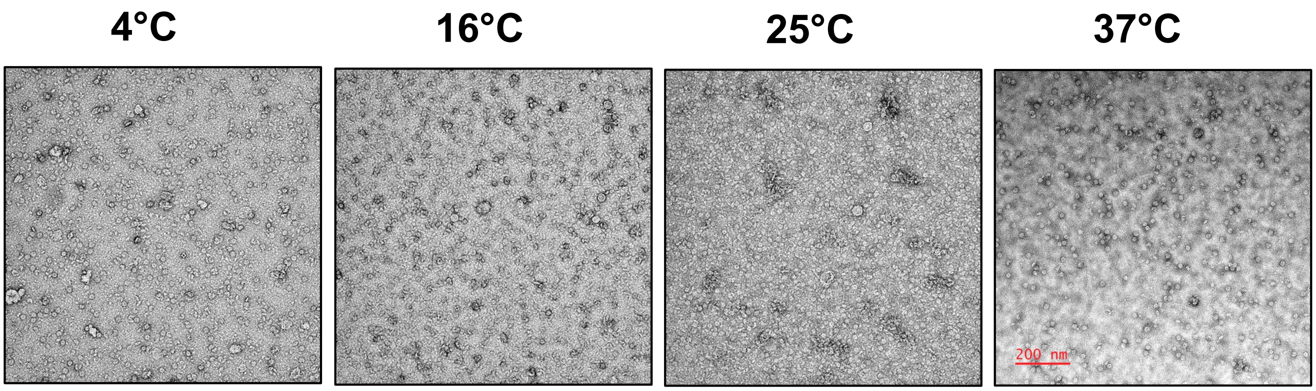


**Figure S1. Portal formation at different temperatures.** Electron micrographs of negatively stained portals produced at 4 °C, 16 °C, 25 °C and 37 °C. Scale bar = 200 nm.

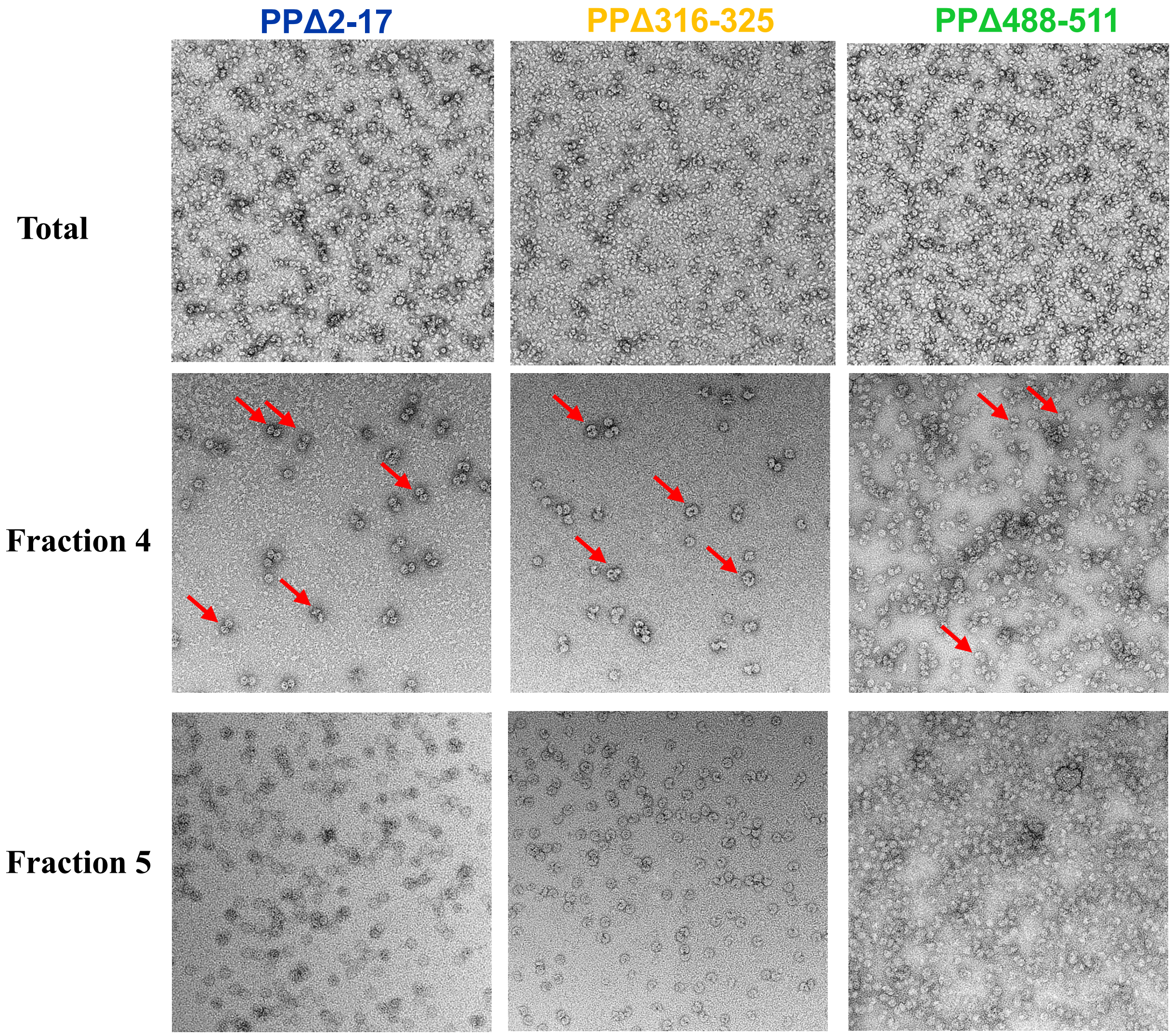


**Figure S2. Negative stain EM of portal deletions.** Portal deletions PPΔ2-17, PPΔ316-325 and PPΔ488-511 affinity purified (Top row, Total) and after SEC separation into fractions 4 and 5. Representatives of doublets are indicated by the red arrows in fraction 4.

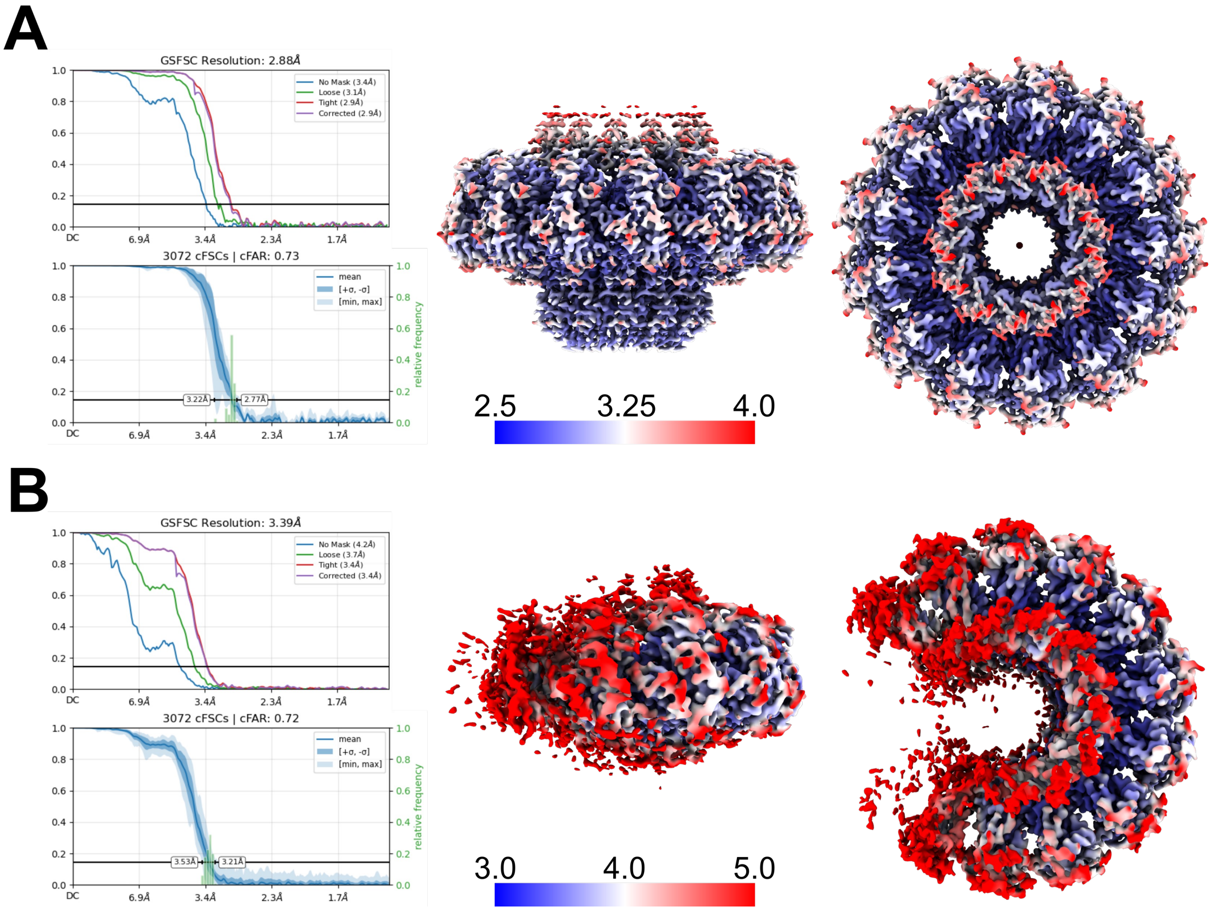


**Figure S3. Reconstruction of portals.** Fourier Shell Correlation (FSC) curves (left) and local resolution maps (right) of the tridecamer **(A)** and unclosed undecamer **(B)** reconstructions from the PPΔ316-325 deletion. The local resolution maps are colored by resolution (in Å) according to the color bar.
